# Widespread deviant patterns of heterozygosity in whole-genome sequencing due to autopolyploidy, repeated elements, and duplication

**DOI:** 10.1101/2023.07.27.550877

**Authors:** Xavier Dallaire, Raphael Bouchard, Philippe Hénault, Gabriela Ulmo-Diaz, Eric Normandeau, Claire Mérot, Louis Bernatchez, Jean-Sébastien Moore

**Affiliations:** Institut de biologie intégrative et des systèmes, Université Laval, Québec, Canada; Centre d’Études Nordiques, Université Laval, Québec, Canada; Ressources Aquatique Québec, Canada; CNRS, UMR 6553 ECOBIO, Université de Rennes, France

**Author notes:** Corresponding author: Xavier Dallaire.

**Keywords:** heterozygosity, salmonid, whole-genome sequencing, paralog, autopolyploid, repetitive DNA

## Abstract

Most population genomic tools rely on accurate SNP calling and filtering to meet their underlying assumptions. However, genomic complexity, due to structural variants, paralogous sequences and repetitive elements, presents significant challenges in assembling contiguous reference genomes. Consequently, short-read resequencing studies can encounter mismapping issues, leading to SNPs that deviate from Mendelian expected patterns of heterozygosity and allelic ratio. In this study, we employed the ngsParalog software to identify such deviant SNPs in whole-genome sequencing data from four species: Arctic Char (*Salvelinus alpinus*), Lake Whitefish (*Coregonus clupeaformis*), Atlantic Salmon (*Salmo salar*), and the American Eel (*Anguilla rostrata*), with low (2X) to intermediate (6X) coverage. The analyses revealed that deviant SNPs accounted for up to 62% of all SNPs in salmonid datasets and approximately 10% in the American Eel dataset. These deviant SNPs were particularly concentrated within repetitive elements and genomic regions that had recently undergone rediploidization in salmonids. Additionally, narrow peaks of elevated coverage were ubiquitous along all four reference genomes, encompassed most deviant SNPs and could be partially attributed to transposons and tandem repeats. Including these deviant SNPs in genomic analyses led to highly distorted site frequency spectra, apparent homogenization of populations and underestimating pairwise F_ST_ values. Considering the widespread occurrence of deviant SNPs arising from a variety of source, their important impact in estimating population parameters, and the availability of effective tools to identify them, we propose that excluding deviant-SNPs from WGS datasets is required to improve genomic inferences for a wide range of taxa and sequencing depths.

Significance: Genomes can be very repetitive and hard to assemble into a reference, which can lead to biases when genotyping genetic markers in complex genomic regions. Here, we draw attention to this issue in various whole-genome datasets and validate a method to identify problematic SNPs at low coverage. We also explore processes creating such SNPs and their consequences on common population genomics analyses.

## Introduction

Single nucleotide polymorphisms (SNPs) are now the most commonly used genetic markers in the field of genomics due to their widespread distribution within genomes, rich information content, and their detectability using a diverse range of sequencing technologies (e.g. RAD sequencing, Andrews et al. 2016; whole-genome sequencing, Fuentes-Pardo & Ruzzante 2017; low-coverage WGS, Therkildsen & Palumbi 2017). Their utilization, however, relies on several assumptions, such as their biallelic nature, adherence to Mendelian inheritance, and, in some cases, independent segregation (absence of linkage disequilibrium). Multiple studies highlight the ongoing need to validate these assumptions (Chen et al. 2014; Jaegle et al. 2023; Hurles 2002), especially in complex genomes such as those that experienced polyploidization.

Polyploidization is the process by which a complete set of chromosomes is multiplied within a single nucleus and then passed on to progenies (Todesco et al. 2020; Zhang et al. 2019). It can happen following hybridization of different species (allopolyploidy), in which case pairs of chromosomes descended from each species conserve preferential bivalent pairing (Cifuentes et al. 2010; Mason & Wendel 2020). In other cases, polyploidy can result from whole-genome duplication (WGD) events within a single genome (autopolyploidy), which creates groups of chromosomes in multivalent recombining pairing during meiosis. This tetrasomic inheritance can last until mutations create enough sequence divergence to reestablish bivalent pairing in a now doubled number of chromosomes, a process known as rediploidization (Lien et al. 2016; Ohno 1970; Weiss & Maluszynska 2004).

Due to a recent WGD in their common ancestor around 88–103 Mya, salmonids have been central to the study of autopolyploidy in animals (Macqueen & Johnston 2014). This WGD is the fourth to occur along the lineages ancestral to salmonids, after two WGDs in early vertebrates (Simakov et al. 2020) and another one in the ancestor of all teleost fishes, around 320-350 Mya (Glasauer & Neuhauss 2014; Jaillon et al. 2004). Following Ss4R, rediploidization occurred at different times across duplicated regions, leading to either ancestral (AORe) or lineage-specific (LORe) ohnolog resolution (Robertson et al. 2017). This is apparent when comparing levels of sequence divergence between pairs of syntenic ohnologs in salmonid assemblies. These ohnologs display from 85% to nearly 100% nucleotide identity (Gundappa et al. 2022; Lien et al. 2016; Mérot et al. 2023; Smith et al. 2022).

An additional source of complexity in salmonid genomes is their relatively high content in transposable elements (TE), which can make up 50-60% of assembled reference sequences (Minkley 2018). In fact, the correspondence in the timing of Ss4R and the proliferation of transposable elements in salmonid genomes suggests that the WGD event may have disrupted TE regulation processes (Lien et al., 2016). Similar to what is observed in hybrids (Hénault et al. 2023; Laporte et al. 2019), TE expansion may in turn have contributed to the sequence divergence and chromosomal rearrangements that enabled rediploidization (Lien et al. 2016). In ray-finned fishes, the abundance of TEs is also closely correlated to genome size (Chalopin et al. 2015; Gao et al. 2016). Given the large genome size of salmonids (2.5-3.0 Gb), such abundance of TE, in combination with high sequence identity between pairs of recently rediploidized chromosomal regions, have considerably hindered efforts to assemble quality reference genomes in salmonids (Lien et al. 2016; Smith et al. 2022).

Due to their socio-economic importance and their scientific relevance for the investigation of many evolutionary and ecological processes (e.g. local adaptation and speciation), salmonids species have been extensively studied using short-read resequencing technologies like RAD-seq (Elmer 2016). By analyzing data from salmonids and other taxa derived from an ancestral WGD, it has become apparent that collapsed assemblies and under-splitting can bias the genotyping of SNPs on highly similar duplicated loci when they are considered as a single region (Harvey et al. 2015). When overlooked, these biases can have an impact on biological and management interpretations derived from genomic data (O’Leary et al. 2018), for example by creating a false signal of differentiation (Larson et al. 2021). SNPs on paralogous sequences or other collapsed loci that are not the result of gene duplication have been identified using a variety of methods, such as the identification of apparent heterozygotes in haploid samples (Hecht et al. 2013; Sánchez et al. 2009) or from their increased depth of coverage compared to single-copy loci (Davey et al. 2013; Dou et al. 2012). McKinney et al. (2017) proposed a method named HDplot that relies on excess of heterozygotes (H) or deviation from the expected 1:1 allelic ratio in heterozygotes (D) to identify SNPs that do not conform to expected patterns of inheritance. We hereafter refer to such variants as “deviant” SNPs, as opposed to “canonical” SNPs that conform to expected patterns of heterozygosity and allelic ration for non-duplicated loci (after Karunarathne et al., 2022).

In Chinook Salmon (*Onchorynchus tshawytscha*; McKinney et al., 2017), deviant SNPs identified by the HDplot method (17% of all SNPs) predominantly matched paralogs identified by haploid mapping (McKinney et al. 2016) and were especially dense in chromosome arms with ongoing residual tetrasomy, suggesting that paralogy stemming from the ancestral WGD is the main source of deviant SNPs in salmonids. This method has since been applied to other salmonids (*O. mykiss* : 22% of deviants, Fraik et al. 2021; *O. kisutch* : Xuereb et al. 2022; *Salvelinus alpinus* : 24%, Dallaire et al. 2021) as well as other taxonomic groups including polyploid trees (e.g. *Pinus cembra*: 85% of deviants, (Rellstab et al. 2019), crustaceans (e.g. *Homarus americanus* : 22 - 40% of deviants, (Dorant et al. 2020, 2022), and ranids (e.g. *Rana luteiventris* : 16% of deviants, Cayuela et al. 2022). In the last two examples, the HDplot method was adapted to identify and genotype Copy Number Variants (CNV) suspected to be associated to transposable elements (Cayuela et al. 2021; Dorant et al. 2020). Altogether, these recent studies highlight the fact that deviant SNPs can also be found in significant proportions in dataset from non-polyploid species.

In recent years, low-coverage whole-genome sequencing (lcWGS) has emerged as an alternative to reduced representation sequencing for population genomic studies in non-model species (Therkildsen & Palumbi 2017). This cost-effective method offers the opportunity for unprecedented sample sizes of whole-genome sequences by reducing the per sample depth of coverage to as low as 0.1X. At low and medium coverages (under 10X), the lack of confidence around individual genotypes can be circumvented by using a growing variety of tools that explicitly account for genotype uncertainty (Lou & Therkildsen, 2021), such as the Genotype Likelihood (GL) framework implemented in ANGSD (Kornelissen et al. 2014).

To assess the prevalence of deviant SNPs in low to intermediate coverage WGS data and identify the main genomic features responsible for their appearance, we applied a common pipeline to new datasets (around 2X of coverage) from two salmonid species and one anguillid and reanalyzed data on Atlantic Salmon (*Salmo salar*; 6X) published in Bertolotti et al. 2020. We first used HDplot to classify deviant SNPs as it is the most widely used method currently, but we also compared its output to that of ngsParalog (Linderoth, 2018). The latter method is not as widely used but presents some potential advantages such as the use of a likelihood approach to avoid low-confidence genotype calling under 10X of coverage. We then mapped canonical and deviant SNPs and compared their relative distribution particularly in relation with peaks of elevated coverage, repetitive regions and regions that experienced delayed rediploidization in salmonids. Finally, we compared the results of common population genomics statistics and analyses before and after filtering for deviant SNPs to assess the impacts of these filters on inferences that are commonly drawn from WGS datasets.

## Results

### Identification and validation of paralog SNPs at low coverage

SNPs flagged as deviant by *ngsParalog* represent a different proportion across datasets but they show consistent characteristics such as low F_IS_ and high coverage. After subsampling, we obtained 5 datasets with an average depth of coverage around 1.5X and a mode between 1.82X and 1.97X (Table 1; see Figure S1 for distribution of depth). We identified SNPs using ANGSD and calculated the likelihood of reads being misaligned at the positions of those SNPs using ngsParalog (p < 0.001). While deviant SNPs were in majority in both Lake Whitefish datasets (49.9% and 61.8%) and in Arctic Char (62.3%), they represented 22.6% of the SNPs in Atlantic Salmon, and 10.6% in the American Eel (Table 1). Nearly all deviant SNPs had F_IS_ under -0.05, while canonical SNPs had a FIS distribution centered around 0 (Figure 1A), except for the Arctic Char dataset where strong population structure (max pairwise F_ST_ = 0.45) created a deficit of heterozygotes (positive F_IS_) putatively due to a Wahlund effect (Wahlund 1928). In datasets with depths of coverage around 1.5X, the average coverage was consistently higher at the positions of deviant SNPs than canonical SNPs (Figure 1D).

**Table 1:** Summary of datasets reanalyzed in this study. See main text for the treatment of each dataset.

| Dataset | Family | GenBank<br>assembly<br>accession<br>(Contig N50) | Origin | Number of<br>sampling sites | Sample size | Sequencer | Original<br>depth of<br>coverage by<br>sample;<br>mean (SD) | Mode of depth in<br>subsampled data | Number of<br>SNP | Number of<br>deviant SNP | % of deviant<br>SNPs |
| --- | --- | --- | --- | --- | --- | --- | --- | --- | --- | --- | --- |
| American Eel<br>(*Anguilla rostrata*) | *Anguillidae* | GCA_018555375.2<br>(N50 = 5.02 Mb) | East coast of<br>North and Central<br>America | 21 | 460 | NovaSeq<br>6000 S4<br>PE150 | 3.68 (0.9) | 1.97 | 16,379,954 | 1,729,598 | 10.60% |
| Arctic Char<br>(*Salvelinus alpinus*) | *Salmonidae* | GCA_002910315.2<br>(N50 = 55 Kb) | Arctic Canada | 16 | 520 |  | 1.65 (0.51) | 1.88 | 5,408,454 | 3,371,454 | 62.30% |
| Lake Whitefish<br>(*Coregonus clupeaformis*) | *Salmonidae* | GCA_018398675.1<br>(N50 = 5.73 Mb) | Great Slave Lake,<br>Canada | 9 | 298 |  | 1.48 (0.34) | 1.63 | 4,491,496 | 2,777,770 | 61.80% |
|  |  |  | James Bay,<br>Canada | 13 | 470 |  | 1.71 (1.90) | 1.88 | 7,318,227 | 3,651,681 | 49.90% |
| Atlantic Salmon<br>(*Salmo salar*) | *Salmonidae* | GCA_905237065.2<br>(N50 = 28.06 Mb) | Coastal Norway | 55 | 126 | HiSeq 2500<br>PE125 | 7.46 (1.72) | 1.82 | 4,392,892 | 990,766 | 22.60% |
|  |  |  |  |  | 228 | HiSeq 3000<br>PE100 | 4.79 (0.75) |  |  |  |  |

**Fig. 1:**
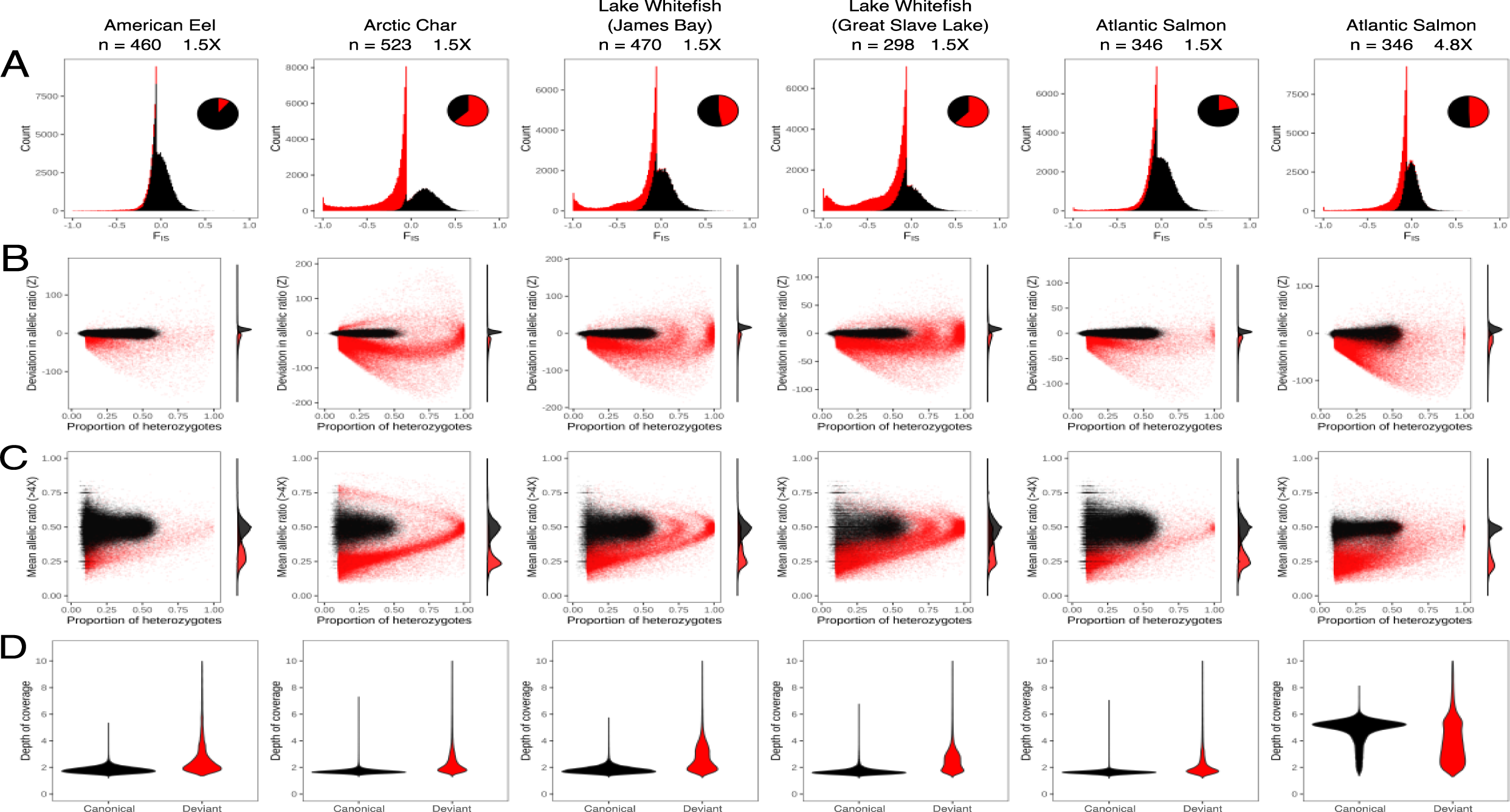
All investigated datasets harbor SNPs in deviation of Hardy-Weinberg Equilibrium and allelic ratio. Summary of canonical (black) and deviant (red) SNPs as categorized by ngsParalog (p < 0.001) for 100,000 randomly selected SNPs in the American Eel, Arctic Char, Lake Whitefish (James Bay and Great Slave Lake), and Atlantic Salmon (1.5X and 4.8X coverage) datasets. A) Histograms of F_IS_ with an inset pie chart showing the proportion of SNPs by category. B) HDplot showing proportion of heterozygotes in relation to deviation in allelic ratio (Z-score). C) Proportion of heterozygotes in relation to mean allelic ratio in heterozygous samples with at least 4X coverage. D) Distribution of depth of coverage by SNP category. The y-axis was restricted to depth under 10X for clarity, but deviant SNPs had maximum depths that greatly exceeded 10X.

The characterization of deviant SNPs showed high consistency between the different methods employed. When visualizing SNPs using HDplot (McKinney, 2017), we observed that the vast majority of SNPs categorized as deviant by *ngsParalog* either had a high proportion of heterozygotes or deviated from the 1:1 expected allelic ratio in heterozygotes (Z further from zero than canonical SNPs) (Figure 1B-C). Among SNPs in excess of heterozygotes (F_IS_ < 0 and p < 0.05 for Hardy-Weinberg Equilibrium test in ANGSD), most were categorized as deviant by ngsParalog: 96.9 to 99.6% in the Lake Whitefish and Arctic Char datasets, 89.7% in the American Eel, and 78.3% in the Atlantic Salmon. For SNPs with lower MAF, the distribution of the average allelic ratio (in heterozygotes with more than 4X of coverage) for deviant SNPs had an obvious mode around 0.25 (1:3) and smaller one at 0.75 (3:1). For canonical SNPs, the average allelic ratio was centered on 0.50 (1:1).

### Prevalence of paralog SNPs at different depths of coverage

We inferred the presence of deviant SNPs in all datasets when depths of coverage ranged from low (1X) to intermediate (4.8X). We categorized SNPs from the Atlantic Salmon dataset before subsampling (4.8X) using *ngsParalog*, then compared the list of SNPs retained in subsampled datasets at decreasing depths of coverage (Figure 2). First, 49.8% of the SNPs (a total of 4.52 millions) were characterized as deviants in the original dataset sequenced at intermediate coverage (4.8X). The number of canonical SNPs and the depth of coverage had a plateau-like relation: it was nearly stable between 4.8X and 3X (93.3% of SNPs left), decreasing slightly between 3X and 1.5X (69.2% of SNPs left), and nearly all canonical SNPs were lost between 1.5X and 1X (less than 0.2% of SNPs left). On the other hand, the number of deviant SNPs (categorized based on the original dataset) did not reach a plateau between 1X and 4.8X, as the number of deviant SNPs was almost directly proportional to the average depth of coverage (Pearson’s R^2^ = 0.98).

**Fig. 2:**
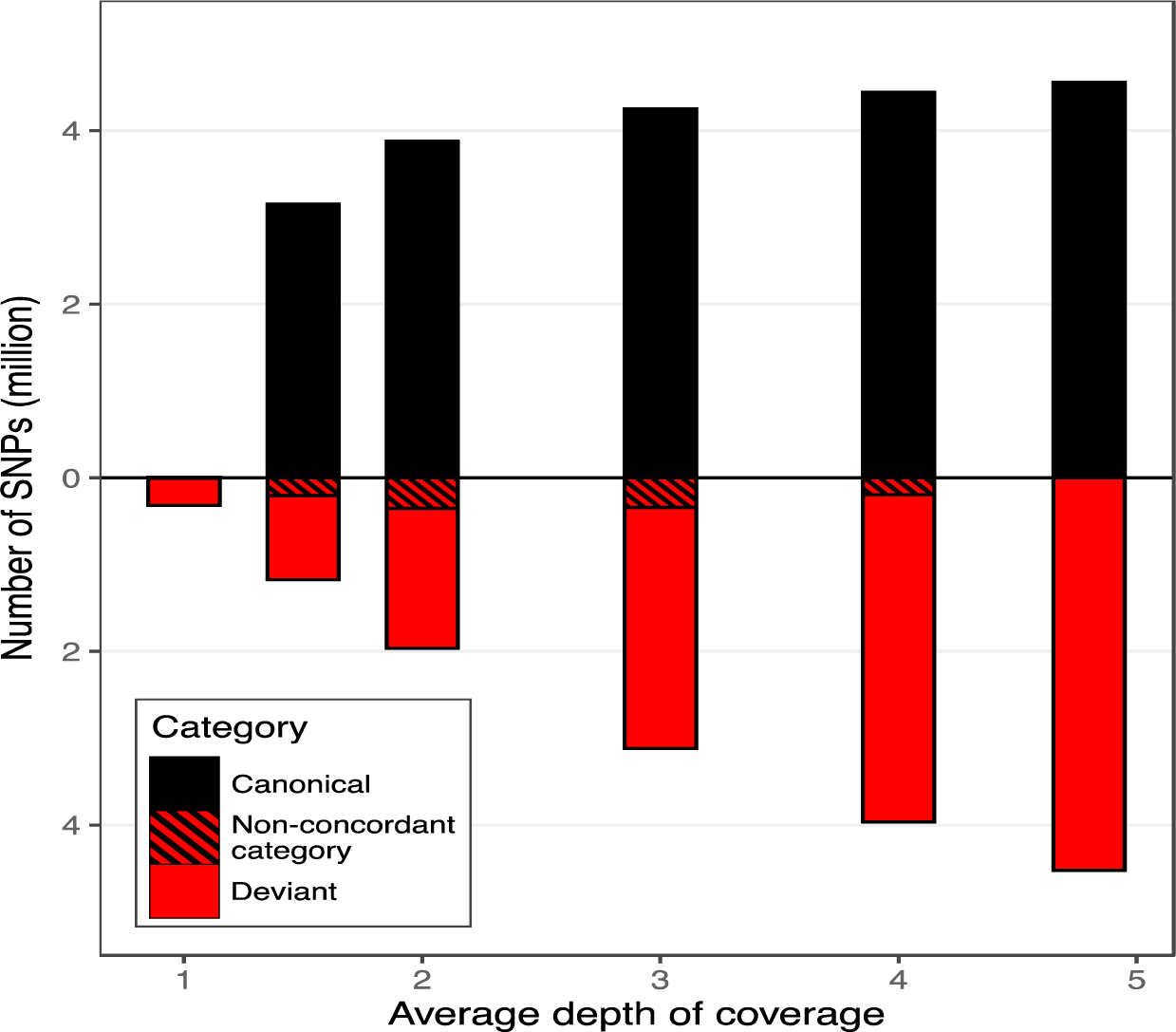
Deviant SNPs are found in low to intermediate coverage datasets. Number of canonical (black, above) and deviant (red, below) SNPs as categorized by ngsParalog in the 4.8X Atlantic Salmon dataset. Deviant SNPs categorized as canonical in the subsampled datasets are represented by the hatched portion of bars, but canonical SNPs categorized as deviant were too rare to be visualized. SNPs absent from the 4.8X dataset (less than 1.5% of all SNPs) were not shown.

We repeated the ngsParalog analysis on the subsampled data and compared the categorization of SNPs found in the 4.8X dataset. Assuming the categories derived from the 4.8X dataset were closer to reality, subsampling the data led to increasing rates of deviant SNPs categorized as canonical (putative false positives) that reached 17.9% of deviant SNPs (5.9% of all SNPs) in the 1.5X dataset. However, canonical SNPs categorized as deviant (putative false negative) were much rarer and peaked at 0.4% of canonical SNPs (0.2% of all SNPs) in the 4X dataset before decreasing in lower coverage datasets.

### Distribution of deviant SNPs in the genome

The distribution of deviant SNPs was at least partly consistent with the hypothesis that they are caused by ancient polyploidization in salmonids. In all three salmonid species, the density of canonical SNPs decreased as the percentage of identity increased between the self-syntenic blocks (ohnolog pairs of sequence resulting from ancient whole-genome duplication), while the density of deviant SNPs increased (Figure 3, Table 2). In Arctic Char, Lake Whitefish, and Atlantic Salmon respectively, 7.9%, 28.6%, and 30.5% of the deviant SNPs were found in self-syntenic blocks with an identity above 95%, while 3.9%, 9.3%, and 12.7% of canonical SNPs were found in those same blocks.

**Fig. 3:**
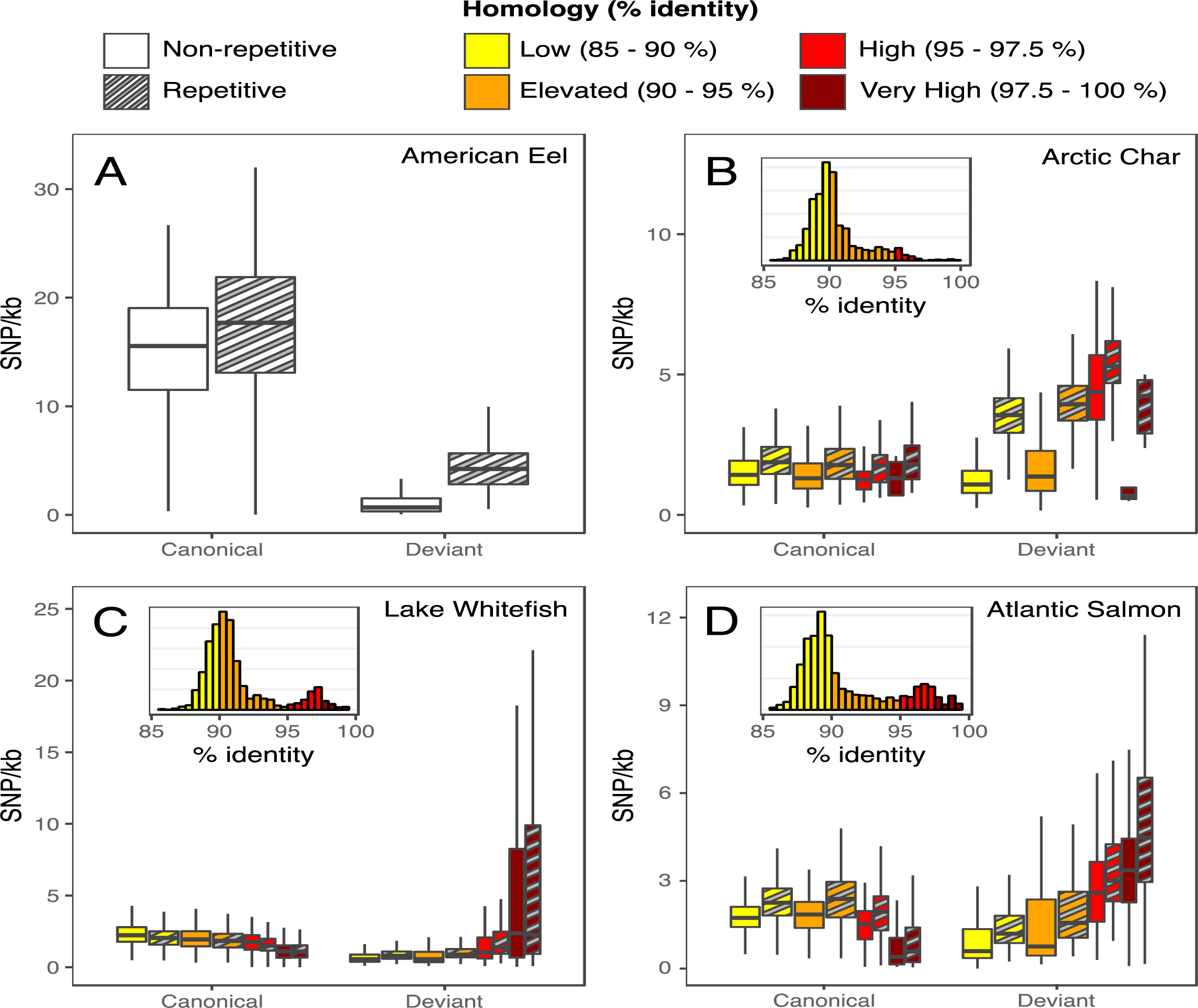
Deviant SNPs are more common in repetitive DNA and recently rediploidized regions. Density of canonical and deviant SNPs (by kb) in 1Mb non-overlapping windows for the A) American Eel, B) Arctic Char, C) James Bay Lake Whitefish, and D) 4.8X Atlantic Salmon datasets. SNPs are split based on whether they were found in repetitive regions identified by Repeat Masker (hatched) or not (plain boxplot). Windows are categorized based on their percentage of identity with their ohnolog, color coded from yellow to dark red. For salmonid datasets, the inset histograms show the relative frequency of percentage of identity in windows. Only base pairs with an average depth above 0.75X were considered for the calculation of SNP densities.

**Table 2:** Parameters of negative binomial models for the rate of canonical or deviant SNPs in 1Mb non-overlapping windows, in relation to non-repetitive and repetitive sequences. The models for salmonids also include the standardized percentage of identity between the window and its ohnolog. Estimates of the fixed effects (β), as well as their 95% confidence intervals (CI), are shown.

| Dataset | Parameter | Canonical SNPs<br>$\beta$ [95% CI] | Deviant SNPs<br>$\beta$ [95% CI] |
| --- | --- | --- | --- |
| American Eel | Intercept | -4.21 [-4.24, -4.18] | -6.58 [-6.63, -6.52] |
|  | Repetitive | 0.14 [0.09, 0.18] | 1.25 [1.16, 1.32] |
| Arctic Char | Intercept | -6.51 [-6.53, -6.48] | -6.44 [-6.46, -6.41] |
|  | Repetitive | 0.25 [0.22, 0.28] | 0.90 [0.86, 0.93] |
|  | Identity | -0.01 [-0.02, 0.01] | 0.22 [0.20, 0.24] |
| Lake Whitefish<br>(James Bay) | Intercept | -6.19 [-6.20, -6.17] | -6.78 [-6.81, -6.74] |
|  | Repetitive | -0.08 [-0.11, -0.06] | 0.09 [0.04, 0.14] |
|  | Identity | -0.11 [-0.13, -0.10] | 0.36 [0.34, 0.38] |
| Atlantic Salmon<br>(4.8X) | Intercept | -6.40 [-6.42, -6.38] | -6.53 [-6.56, -6.49] |
|  | Repetitive | 0.27 [0.24, 0.29] | 0.32 [0.28, 0.37] |
|  | Identity | -0.14 [-0.15, -0.12] | 0.36 [0.34, 0.38] |

The distribution of deviant SNPs was also associated with the presence of repetitive regions (i.e. interspersed and tandem repeats). For American Eel, Arctic Char, and Atlantic Salmon, both canonical and deviant SNPs were found in higher density in repetitive regions than in non-repetitive regions, but the effect was strongest for deviant SNPs in American Eel and Arctic Char (Figure 3, Table 2). In Lake Whitefish, canonical SNPs were less dense in repetitive regions, while deviant SNPs were denser.

The association between deviant SNPs and repeated elements was evident when looking at peaks of elevated coverage found in all datasets and that varied in size in the order of tens to hundreds of bp (Table 3). While these high-coverage regions covered a relatively small portion of each genome, deviant SNPs were clearly over-represented in those peaks of coverage (χ^2^ > 493,507, ddf = 1) while canonical SNPs were under-represented (χ^2^ > 3,230). We explored the interaction between depth of coverage, deviant SNPs, and repetitive elements (Fig. 4A). Depending on the dataset, the peaks of elevated coverage were enriched in certain types of repetitive elements, namely long terminal repeat (LTR) retrotransposons, satellites, simple repeats, and low complexity regions, as defined by RepeatMasker (Figure 4B, Table S1). This enrichment was especially strong and extended to most repeat types in the American Eel dataset.

**Table 3:** Characteristics of four datasets regarding the sufficiently covered part of the genome (depth of coverage above 0.75X) and peaks of elevated coverage (above 3X in American Eel, Arctic Char, and Lake Whitefish; above 8X in Atlantic Salmon). Percentage of canonical and deviant SNPs (as categorized by ngsParalog) inside peaks of coverage are shown.

| Species | % of the genome sufficiently covered | Median peak width (bp) | 95th quantile of peak width (bp) | % of sequenced genome in peaks | % of canonical SNPs in peaks | % of deviant SNPs in peaks |
| --- | --- | --- | --- | --- | --- | --- |
| American Eel | 92.50% | 30 | 329 | 2.10% | 1.00% | 61.70% |
| Arctic Char | 83.40% | 76 | 398 | 3.20% | 1.30% | 59.70% |
| Lake Whitefish (James Bay) | 83.30% | 39 | 821 | 4.60% | 1.90% | 64.40% |
| Atlantic Salmon (4.8X) | 90.20% | 65 | 457 | 1.80% | 0.40% | 27.60% |

**Fig. 4:**
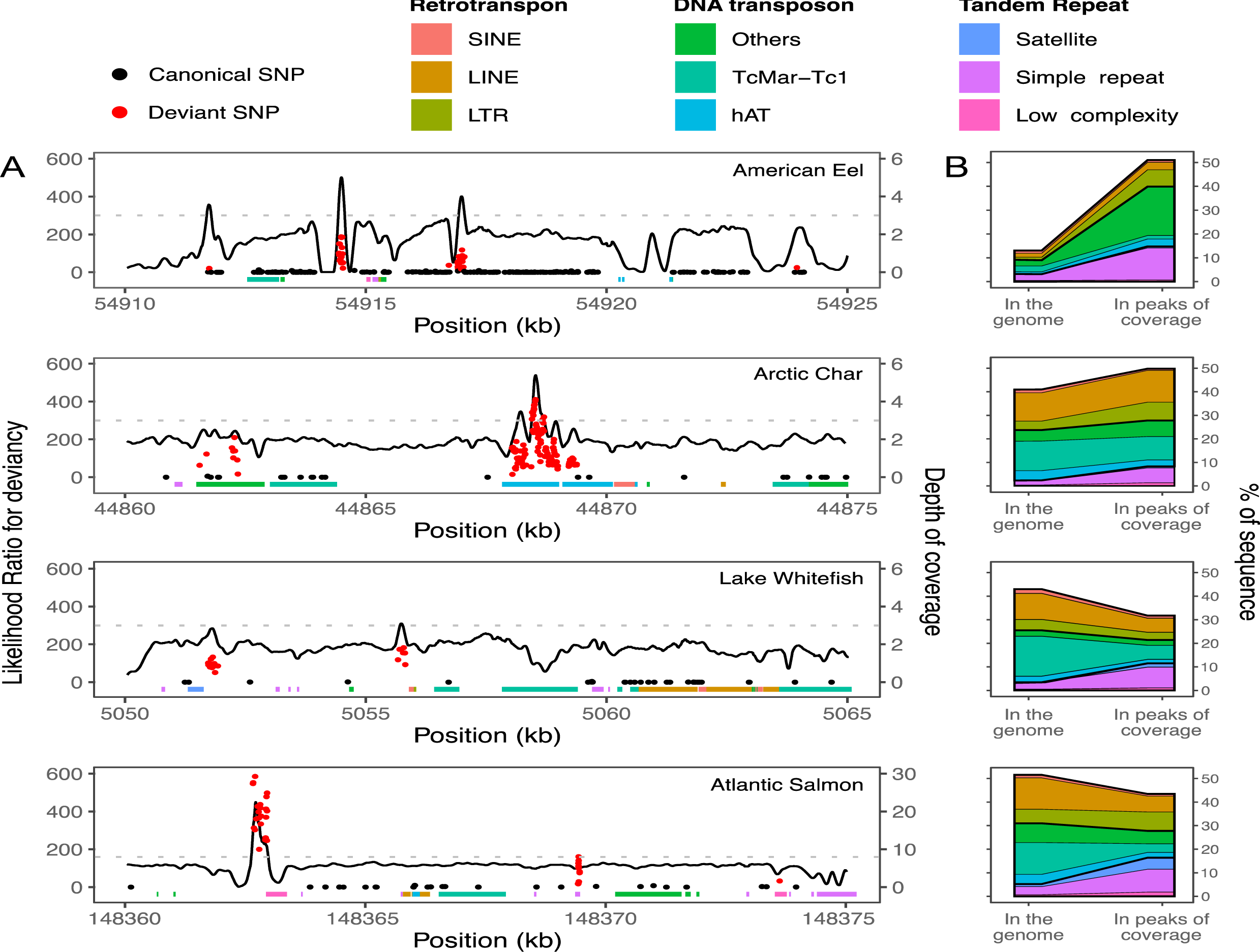
Peaks of elevated coverage are enriched in both deviant SNPs and certain classes of repetitive DNA elements. A) Depth of coverage in 15 kb windows on the first chromosome of the American Eel, Arctic Char, Lake Whitefish, and Atlantic Salmon (from top to bottom; the first chromosome was arbitrarily selected and is shown here as a representative illustration of patterns observed over the entire genome). The position of canonical (black) and deviant (red) SNPs are marked as points according to their likelihood ratio of being in a mismapped region, according to ngsParalog. The extent of repetitive elements is indicated by colored rectangles at the bottom of the plots, and the depth threshold for peaks of coverage is shown by the light grey dashed line. B) Proportion of sequence covered of the most frequent clades of transposable elements and other repeat types in the sufficiently covered portion of the genome (left) compared to peaks of elevated coverage between 20 and 1,000 bp (right).

### Effects of deviant SNPs on population genomics analysis

Including deviant SNPs or excluding them had a strong impact on population genomic statistics, such as shared polymorphism and genetic differentiation. When comparing SNPs found for Whitefish in James Bay (JB; 7,318,277 SNPs) and in Great Slave Lake (GSL; 4,491,496 SNPs), we found 4,037,742 SNPs at common positions, 3,714,622 of which had the same alternative alleles. Those common polymorphisms thus represented 82.7% of those found in GSL and deviant SNPs were found in similar proportions in the common list (65.8%) and in GSL (67.0%). However, 3.7% of the common SNPs were categorized as deviant only in the GSL dataset, and 2.7% were deviant only according to the JB dataset (Figure S2A). Common SNPs categorized as deviant in both datasets showed highly correlated MAF (R^2^ = 0.90, Figure S2B) and F_IS_ (R^2^ = 0.90, Figure S2C), compared to other SNPs (canonical: R^2^ = 0.63 and 0.07; SNPs with non-concordant categories: R^2^ = 0.56 and -0.31).

In the Arctic Char dataset, pairwise F_ST_ values estimated with all SNPs and only canonical SNPs were highly correlated (R^2^ = 0.88), but estimates were 2.02 times higher when using only canonical SNPs (Figure 5A). When inspecting two-dimension site frequency spectra between sampling sites with different levels of divergence, we found that deviant SNPs were consistently located along the diagonal and concentrated at low (< 0.1) and high (0.5) MAF (Figure 5B-C). To summarize, failure to filter out deviant SNPs leads to underestimating genetic differentiation indices between populations by increasing the number of shared polymorphisms at similar frequencies.

**Fig. 5:**
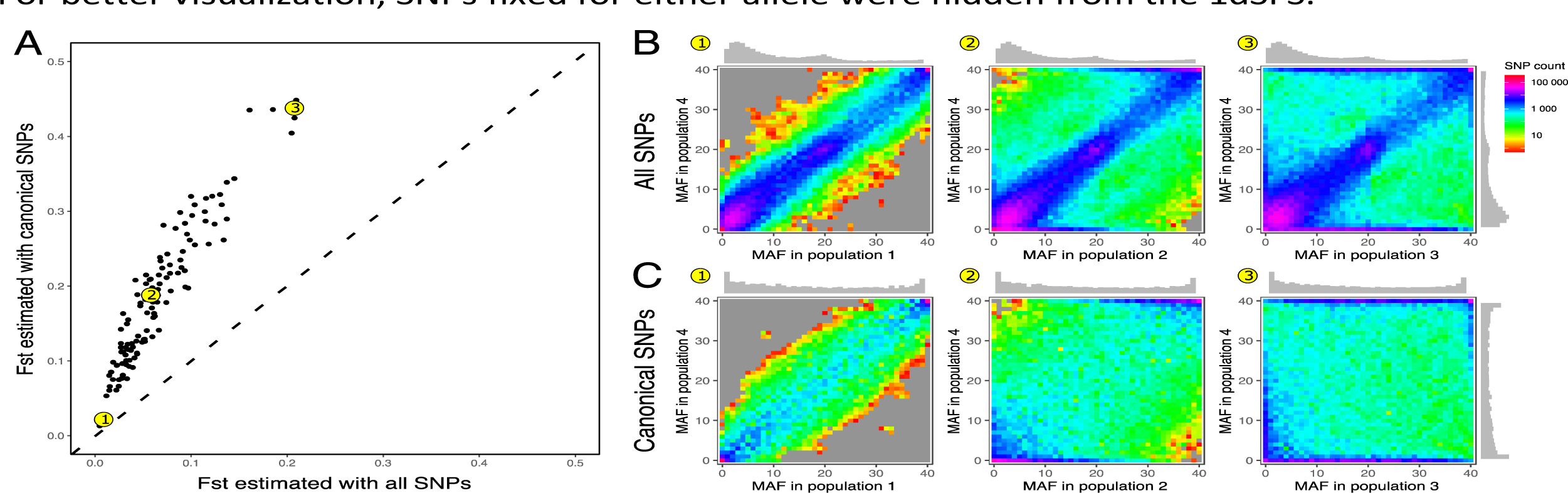
Population genetic differentiation is underestimated when deviant SNPs are not removed from the dataset. A) Relationship between pairwise Fst estimated using all SNPs in the Arctic Char dataset and only SNPs categorized as canonical by ngsParalog. Three pairs of populations are highlighted in yellow, corresponding to population 1 (low Fst), 2 (medium Fst), and 3 (high Fst) paired with a fourth common population. Unfolded two-dimensional site frequency spectra (2dSFS) are shown for these pairs using B) all SNPs and C) only canonical SNPs. One-dimensional SFS for the four highlighted populations are shown along the axes of the 2dSFS. For better visualization, SNPs fixed for either allele were hidden from the 1dSFS.

## Discussion

In this study, we aimed at investigating the prevalence of deviant SNPS in WGS data by applying a common variant calling pipeline to both new and previously published datasets covering four different fish species. We found a significant proportion of SNPs to be in deviation from expected patterns of heterozygosity and allelic ratio in salmonid datasets, and to a lesser extent in the American Eel. We attribute most of these deviant SNPs to collapsed assembled genomic regions, which is frequent in salmonid assemblies because of a recent rediploidization, as well as to repetitive sequences. Considering the widespread occurrence of deviant SNPs arising from a variety of source, their important impact in estimating population parameters, and the availability of effective tools to identify them, we suggest that excluding deviant-SNPs from WGS datasets is required to improve genomic inferences for a wide range of taxa and sequencing depths.

### Detection of paralogs in low-coverage whole-genome sequencing datasets

Identifying SNPs on paralogous or other multi-copy sequences has been a long-lasting challenge in genomics. The HDplot method (McKinney et al. 2017) is now commonly applied to detect such SNPs based on their deviation from Hardy-Weinberg Equilibrium and from the expected allelic ratio in heterozygotes. *ngsParalog* (Linderoth 2018) is an openly available bio-informatic software that uses similar signals to test the hypothesis that the mismapping of reads creates deviant SNPs at specific genomic positions. By using a probabilistic approach that avoids the use of strict genotyping and taking into account the uncertainty inherent to low-coverage SNP calling, *ngsParalog* is reported to be able to detect deviant SNPs in Next Generation Sequencing datasets with coverage as low as 2X (Linderoth, 2018). This tool has been used both on RADseq (Benjamin et al. 2018; Hemstrom et al. 2022; Saglam et al. 2017) and WGS (Márquez et al. 2020; Pope et al. 2023) datasets. However, none of these studies reported the number of SNPs filtered out by this approach.

Here, we applied *ngsParalog* on a variety of datasets for salmonid species where we expected high numbers of deviant SNPs, as well as one non-salmonid fish. By using the HDplot as a visual validation, we observed that deviant SNPs detected by *ngsParalog* presented observed heterozygosities and allelic ratios in perfect accordance with the theoretical and simulated distributions presented in McKinney et al. (2017). Deviant SNPs were very common in the Arctic Char and Whitefish datasets, representing >50% of all SNPs. While we cannot evaluate accurate false positive and negative rates, we can infer by comparing the two Whitefish datasets (n = 298 and 470) that increased sample size led to better defined groups of SNPs (canonical vs deviant) in the HDplot. This is likely also reflected in the detection power for deviant SNPs when using ngsParalog, as this method uses diagnostic factors similar to those used with the HDplot method. Since these methods depend on accurate measurements from heterozygous samples, we suggest that deviant SNPs identification should be done using the whole dataset rather than at population level in order to maximize sample size and thus power.

The subsampled Atlantic Salmon dataset (1.5X) had a much lower proportion of deviant SNPs than other salmonid datasets, but this was not the case in the original higher coverage data (4.8X). In fact, subsampling the Atlantic Salmon dataset mainly led to the loss of deviant SNPs, while canonical SNPs were mostly conserved. However, it is difficult to predict how an increased sequencing effort would affect the already high proportions of deviant SNPs in the Arctic Char and Lake Whitefish datasets, because those differ in many ways from the Atlantic Salmon data. First, we used a recent version of the Atlantic Salmon reference genome (Ssal_v3.1), assembled with more resources and of better overall quality than the *Salvelinus sp.* and *Coregonus clupeaformis* references (Table 1). Second, while all other datasets were generated by a common protocol using Nextera librairies and Illumina NovaSeq 6000 S4 (paired-end reads of 150 bp), the Atlantic Salmon data was pieced together from multiple batches on Illumina HiSeq (read length of 100 or 125 bp). Since deviant SNPs are above all caused by mismapping on the reference genome as shown above, these differences might influence the main source of deviant SNPs in the Atlantic Salmon data. This is exemplified by the fact that paralogs were not as predominantly associated with peaks of elevated coverage as in the other datasets. Nevertheless, our analyses at different depths of coverage in Atlantic Salmon strongly suggest that high proportions of deviant SNPs are not exclusive to very low coverage datasets.

For organisms without recent polyploidization (i.e. not expected to produce datasets harboring high levels of paralogs), like the American Eel included here, common filters include setting a maximum depth of coverage per SNPs (e.g. four times the average depth). This should avoid SNP calling in collapsed regions where the alignment of reads from multiple loci leads to a localized increase in coverage. However, in the American Eel data, canonical and deviant SNPs had largely overlapping depth distributions, and only 3.2% of the deviant SNPs had extreme values for depth of coverage (>8X). Another common approach is to remove SNPs with strong excess of heterozygotes, since few biological processes can explain such patterns in the distribution of genotypes. However, we found that we did not have the power to detect such excesses for most deviants in the American Eel datasets as they appeared to be at much smaller MAFs (and thus expected heterozygosities) than in Arctic Char and Lake Whitefish. This might be caused by panmixia in the American Eel (Côté et al. 2013; Ulmo Diaz et al., *in prep*) or by higher copy numbers for deviant SNPs created by repetitive DNA than those arising from residual tetrasomy and delayed rediploidization like in salmonids. Based on these observations, we argue that simpler filtering steps might miss the majority of deviant SNPs in a wide variety of datasets, and that multiple factors (i.e.: excess of heterozygotes, deviation from expected allelic ratio and depth of coverage) should be jointly considered when filtering deviants. As such, we found *ngsParalog* to offer an resource-unintensive and multi-factor program for reliable deviant SNP filtration that could easily be applied to all WGS datasets.

### Distribution of deviant SNPs

Next, we investigated the main processes leading to the presence of deviant SNPs in the analyzed datasets by comparing the distribution of canonical and deviant SNPs along the genome. Our observations were consistent with the prediction that deviant SNPs were more frequent in (1) regions of elevated homology due to delayed rediploidization (LORe) and (2) in repetitive elements. First, LORe regions either have very recently started to differentiate or are still experiencing tetrasomic recombination (Robertson et al. 2017; Waples et al. 2016). This creates important challenges for the linearization of those sequences in reference genomes, which might result in the collapse of both ohnologs into a single consensus sequence, the mismapping of reads, and the creation of deviant SNPs. This is in line with the RADseq data on Chinook Salmon shown in McKinney et al. (2017), where putative paralogs were found almost exclusively in chromosome arms expected to have experienced LORe.

However, we found deviant SNPs to be ubiquitously distributed along all 4 studied genomes. This suggests that delayed rediploidization following a whole-genome duplication is not the only factor at play, since up to half of deviant SNPs in salmonid datasets were distributed outside of LORe regions. Moreover, we also found numerous deviant SNPs in the American Eel, for which the last WGD event is much older than in salmonids. We observed that deviant SNPs were more frequent inside repetitive sequences, and that this effect was especially strong in the American Eel. This supports the idea that interspersed and tandem repeats are another significant source of collapsed assemblies and mismapping. This seemed to happen in narrow peaks of elevated coverage which were extremely dense in deviant SNPs (Figure 4). Those peaks were of similar size or smaller than reads and were most often disproportionally associated to micro- and minisatellites, as well as long-terminal repeat (LTR) transposons. Both transposable elements and tandem repeats, such as satellites, are notoriously challenging to assemble (Sotero-Caio et al. 2017; Tørresen et al. 2019; Treangen & Salzberg 2012), possibly even when using long-read technologies (Liljegren et al. 2016). Tandem repeats are sometimes referred to as genomic “dark matter” in that they are nearly impossible to assemble (Sedlazeck et al. 2018; Weissensteiner et al. 2020) and they hamper the contiguity of assemblies by creating gaps (Peona et al. 2021; Star et al. 2011).

The genome-wide distribution of deviant SNPs we observed here thus apparently arises from a multitude of sources. While some sources of deviant SNPs remain cryptic, we identified processes specific to organisms with recent polyploid ancestors, i.e. the lingering homology between ohnolog pairs of chromosome, as well as some processes common to all organisms, i.e. repetitive elements. These observations suggest that the problems caused by deviant SNPs are not restricted to highly complex or recently duplicated genomes.

### Consequences of deviant SNPs

Our analyses support the idea that deviant SNPs, no matter their origin, should be removed from genomic datasets before proceeding with any population-level analyses. Indeed, we showed that deviant SNPs can create noise or biased signals when quantifying and interpreting the extant of genetic diversity within and between populations. For example, Verdu et al. (2016) reported an overestimation of genetic diversity when paralogs where unaccounted for, as would be expected with the inclusions of numerous SNPs with inflated heterozygosities. Diverged duplicates, i.e. spurious SNP resulting from two collapsed loci fixed for different alleles, would also increase the apparent genetic diversity as they appear polymorphic. An abundance of deviant SNPs in the dataset could even obscure fine-scale population structure, as reported in Atlantic Salmon when comparing F_ST_ with unfiltered and filtered data using PMERGE (Nadukkalam Ravindran et al. 2018). Here, we observed that deviant SNPs displayed shared frequencies, even over genetically diverged groups of individuals (e.g. the two geographically distinct Whitefish datasets or distant Arctic Char populations), leading to distorted joint Sites Frequency Spectra (jSFS). This would create an apparent genetic homogenization of populations, as exemplified by the underestimation of F_ST_ in the Arctic Char dataset before filtering for deviant SNPs.

Apart from F_ST_ estimation, SFS and their joint form are the basis for many other analyses in population genomics, such as neutrality tests (Fu & Li 1993; Tajima 1989), selection scans (Andolfatto 2007; Begun et al. 2007), and demographic inferences (Excoffier et al. 2013; Gutenkunst et al. 2009; Rougeux et al. 2017). Hence, we expect the inclusion of large numbers of SNPs with biased allele frequencies to have more extensive impacts than the underestimation of population differentiation. For example, the enrichment in both low and high frequency sites might result in positive or close to zero values for Tajima’s D, that could be interpreted as spurious signal for balanced selection when coupled with elevated heterozygosity and shared polymorphism across the studied system (Fijarczyk & Babik 2015). It is hard to predict how the disrupting impacts of deviant SNPs scale with their abundance. However, given the relatively low-effort options available to identify and filter such SNPs, the cautionary principle should be applied and they should be removed from all datasets, even when studying species whose genomes contain comparatively fewer repetitive regions.

### Perspectives on best practices

Based on our analysis of multiple WGS datasets, we argue that most if not all NGS datasets would benefit from a rigorous identification of SNPs deviating from expected patterns of heterozygosity and allelic ratios. Indeed, multiple and sometimes cryptic mechanisms can result in the misalignment of short-read sequences. Although our analyses were limited to low and intermediate coverage, we can extrapolate that numerous deviant SNPs could also be found in higher coverage WGS datasets. However, due to budgetary constraints, such datasets are usually characterized by smaller sample sizes, and the resulting decrease in the power to detect deviant SNPs might have obscured the magnitude of the issue until now.

A long-term objective should be to continuously aim for improved genome assemblies, but such concerns fall outside the scope of most resequencing studies, in which the most obvious course of action remains to identify and remove deviant SNPs. This might result in the need to plan for increased sequencing effort when aiming for a specific depth of coverage in complex genomes, as multi-copy loci could gather more reads and decrease the effective coverage for canonical SNPs. Alternatively, some progress is being made to mitigate the problem at the source and improve the design of sequencing libraries. For example, duplex-specific nucleases have been used in library preparation to degrade repetitive sequences, increasing the concentration of single-copy fragments to be sequenced (Matvienko et al., 2013; Shagina et al., 2010; Todesco et al., 2020), thus improving the effective depth of coverage for a similar cost.

It is important to note, however, that the extensive and stringent filtering proposed here still retains millions of high-quality canonical SNPs. We show how appropriate filtration applied on rediploidized and other genomes contribute to make low-coverage whole-genome sequencing a major improvement over reduced-representation sequencing techniques. Indeed, a higher density of variants, coupled with substantial sample size, have the potential to lead to important advancement for a wide array of applications, such as the identification of peaks of differentiation (genomic regions putatively under divergent selection), genome-wide association studies, and gene-environment interactions.

## Methodology

### Sequencing and preprocessing data

To assess the presence of deviant SNPs in studies using low-coverage sequencing, we applied a common detection pipeline to five datasets, four from salmonid species (Arctic Char; Atlantic Salmon; and two datasets from Lake Whitefish, *Coregonus clupeaformis*) and one from a non-salmonid teleost fish (American Eel, *Anguillida rostrata*). The Arctic Char, Lake Whitefish, and American Eel datasets were produced by whole-genome sequencing (Illumina NovaSeq 6000 S4 PE150) on Illumina Nextera libraries made from tissues preserved in Ethanol 95% or RNAlater and extracted by Nucleomag kits (Arctic Char and Lake Whitefish) or the salt-based extraction protocol developed by Aljanabi & Martinez (1997), modified to add a RNase A treatment (Qiagen) (American Eel). The Atlantic Salmon dataset was published in (Bertolotti et al. 2020) and is available on NCBI SRA (accession numbers in Table S2). To limit the potential impacts of transatlantic population structure in this dataset, we downloaded raw sequences from Norwegian samples only, which were sequenced either on Illumina HiSeq 2500 PE125 or HiSeq3000 PE100. Samples on either batch were distributed across the study range on the coast of Norway (Figure S3).

All data were prepared following the pipeline described at https://github.com/enormandeau/wgs_sample_preparation. In brief, raw sequences were trimmed using *fastp* (Chen et al. 2018) and aligned on their respective reference genome (Table 1) with *bwa mem* (minimum alignment quality of 10). Duplicate reads were then removed using *picard*, indels were realigned and overlapping ends of paired reads were clipped. Average per base depth of coverage was then estimated using *samtools depth*. To mitigate batch effects in the Atlantic Salmon dataset, sequences obtained on HiSeq 2500 PE125 were randomly subsampled using *samtools view -s* with a factor of 0.64 to normalize both sequencer batches around an average coverage of 4.8X while maintaining variation in coverage between individuals. Three of these samples (Alta_12_0228, Arga_12_0089, Naus_12_0059) that had initial coverages over 12X were instead subsampled to a target coverage of 4.8X. The normalized (4.8X) Atlantic Salmon dataset was further subsampled to create datasets of decreasing average coverages (4X, 3X, 2X, 1.5X and 1X). Reads from the American Eel data were also randomly subsampled with a factor of 0.5 to reach a similar depth as the Arctic Char and Lake Whitefish datasets.

### Identification and characterization of SNPs

For each dataset, single nucleotide polymorphisms (SNP) were detected using ANGSD v0.931 (Korneliussen et al. 2014) with the GATK genotype likelihood framework (*-GL 2*) and the following parameters. Only reads in a properly mapped pair, with a sequencing quality over 20, and a mapping quality over 30 were considered for SNP calling. Biallelic SNPs were kept when sequenced with a coverage of at least 1X in 75% of all samples and with a minor allele frequency (MAF) above 0.05 *(-doMaf 1 -minMaf 0.05*). Hardy-Weinberg equilibrium was assessed based on the global MAF *(-doHWE 1*) and individual read counts were extracted for each allele *(-doCounts 1 -dumpCounts 4*).

To categorize SNPs as either canonical or deviant, we used the calcLR function in *ngsParalog* (https://github.com/tplinderoth/ngsParalog) that compares the likelihood that reads at the position of SNPs come from either one or multiple copies of a loci. These hypotheses are tested assuming Hardy-Weinberg expectations for non-duplicated loci and using a genotype likelihood framework to account for low-coverage data (Linderoth, 2018). The likelihood ratios of both hypotheses were then compared to a χ^2^ distribution (1 degree of freedom) and SNPs with a p-value (adjusted by applying the Benjamini-Hochberg procedure) under the conservative threshold of 0.001 were considered as deviant. We summed the number of reads for each SNPs to compare depths of coverage between canonical and deviant SNPs.

To validate deviant SNPs, we used an alternative to ngsParalog, consisting in two tests adapted from the HDplot method (McKinney et al. 2017). We first assessed excesses of heterozygotes based on the ANGSD -doHWE 1 output (p < 0.05 and F_IS_ < 0), then computed the deviation from expected allelic ratio in heterozygotes (1:1), following Karunarathne et al. (2022)’s implementation of HDplot. In brief, we used individual read ratios for the alternative allele in heterozygotes to compute a Z-score for each SNP and compared it to a probability density function with a standard deviation of √𝑛, where n is the number of heterozygotes. SNPs displaying an excess of heterozygotes or outside of the 0.025 and 0.975 quantiles for the probability density function were considered deviant.

### Distribution of deviant SNPs in the genome

We ran RepeatMasker v.4.0.8 (Smit et al. 2013) on all four reference genomes to soft-mask transposable elements and other repeats based on the combined DFam (Hubley et al. 2016) and Repbase (Bao et al. 2015) databases for teleost fishes. We masked 45.5% of the *Salvelinus sp.* genome, 52.0% of the Lake Whitefish genome, 49.5% of the Atlantic Salmon genome, and 12.7% of the American Eel genome (Table S3). We then defined “sufficiently covered” regions of the genome by measuring the average depth of coverage of every base pair in each datasets using *samtools depth* and delimiting all segments with an average coverage above 0.75X (i.e the minimum coverage allowing SNP calling according to our ANGSD parameters). In non-overlapping 1Mb windows, we separately counted the number of canonical and deviant SNPs in soft-masked and unmasked sequences, then converted this number in density of SNPs by dividing it by the total length of the sufficiently covered soft-masked or unmasked sequences in the window, respectively.

To estimate the remaining level of homology following the rediploidization of duplicated chromosomes in salmonids, we identified blocks of synteny, i.e. ohnolog pairs of genomic regions descending from the whole-genome duplication. To do so, we hard-masked the repeats identified above then aligned each salmonid reference genome used in this study on itself using *MUMmer v3.23* (Kurtz et al. 2004), implemented in *SyMap v4.2* (Soderlund et al. 2011). We found syntenic blocks (Table S4) concordant with those reported in the original reference for those assemblies (Christensen et al. 2018; Lien et al. 2016; Mérot et al. 2023). We then used *lastz v1.04.15* (Harris 2007) to realign each ohnolog pairs (with arguments *--gfextend --chain --nogapped*), and averaged percentage of identity in non-overlapping windows of 1Mb on all chromosomes. Only windows where *lastz* anchors covered at least 1% of the sequence were kept.

To assess the impact of delayed rediploidization (increased homology) and repeated elements on the risk of mismapping reads and creating deviant SNPs, we created two negative binomial models per dataset, for canonical and deviant SNPs respectively. The number of SNPs in either the soft-masked or unmasked fraction of the window was used as response variable, and the length of the sufficiently covered soft-masked or unmasked fraction of the window was set as an offset to model the density of SNPs rather than the absolute count. The repetitive status (soft-masked or unmasked) and the average percentage of identity in the window were treated as fixed effects for the salmonid datasets. For the American Eel dataset, similar models were built with the repetitive status as the only fixed effect. The negative binomial model was selected after checking for overdispersion of the data in Poisson regressions (c-hat > 1). We checked the goodness-of-fit of the 8 negative binomial models using hanging rootograms (Figure S4).

### Peaks of coverage and transposable elements

To better understand the interaction between deviant SNPs, sequencing depth, and repetitive elements in the reference genome of each species, we catalogued “peaks of coverage” where average depth was elevated compared to the rest of the genome We set the threshold 1.5 times over the mode of depth in each dataset, corresponding to 3X for the Arctic Char, Lake Whitefish (James Bay), and American Eel datasets, and 8X for the Atlantic Salmon (4.8X) dataset. We measured the width of the peaks of coverage, then counted canonical and deviant SNPs occurring inside and outside peaks.

To test the hypothesis that peaks of coverage are enriched in repetitive DNA (which we hypothesized to be one possible cause of peaks of coverage), we used a χ^2^ test to compare the proportion of base pairs covered by different clades of TEs and other repeats in the sufficiently covered portion of the genome and in peaks of coverage.

### Impact of deviant SNPs on population genomics analyses

We compared the list of SNPs in the two Lake Whitefish datasets and their categorization as either canonical or deviant in each dataset. For SNPs common to both datasets, we calculated the Pearson correlation between the MAF and F_IS_ in one dataset and the other. For the Arctic Char dataset, we constructed an ancestral reference genome to polarize alleles. This was done by aligning WGS data from 4 closely related species (Atlantic Salmon; Lake Trout, *Salvelinus namaycush*; Rainbow Trout, *Oncorhynchus mykiss*; and Chinook Salmon, *Oncorhynchus tshawytscha*; SRA accession number in Table S5) on the *Salvelinus sp.* reference genome (ASM291031v2) and using the most common allele as ancestral. We constructed two-dimension site frequency spectra (2dsfs) and calculated pairwise F_ST_ between each population using the argument *-dosaf 1* in ANGSD and the *realsfs fst index*, *print*, and *stats* functions. We repeated this using first the complete list of SNPs, then only the canonical SNPs.

## Acknowledgement

This work was supported by Genome Canada and Genome Quebec as part of “FISHES : Fostering Small-scale Fisheries for Health, Economy and Food Security”. Sampling was made possible by the collaboration of Fisheries and Ocean Canada (Les N Harris, Simon Willey, Ross Tallman, Xinhua Zhu, David Boguski); Ministère de l’Environnement, de la Lutte contre les changements climatiques, de la Faune et des Parcs (Québec; Julien Mainguy); Government of Nunavut; Makivik Corporation (Nunavik); Eeyou Marine Region Wildlife Board (Natacha Louttit, Stephanie Varty); Cree Trappers Association (Sanford Diamond, George Natawapineskum, John Lameboy); and numerous Inuit, Cree, and Dene communities and local fishermen across Canada. Thanks to Anne Beemelmanns, Charles Babin, Bérénice Bougas, Alysse Perreault-Payette, and Gabriel Piette-Lauzière for their help in labwork and coordination. We also want to thank Anne-Marie Dion-Côté for her advice on repeated elements, and Alicia C. Bertolotti, Daniel Macqueen, their co-authors and the Norwegian University of Life Sciences for providing the Atlantic Salmon data. Finally, thank you to the bioinformatics and genomic platform teams at Institut de Biologie Intégrative et des Systèmes (IBIS) for their support.

## Data Accessibility Statement

Upon acceptation of the article, all new genetic data for American Eel, Arctic Char, and Lake Whitefish will be deposited on SRA. Existing data for the Atlantic Salmon was published in Bertolotti et al. 2020 and is available as part of project PRJEB38061 on the European Nucleotide Archive.

The code developed to call SNPs, identify deviant SNPs and compute pairwise F_ST_ will be available upon acceptation of the article on GitHub, as a branch of the pipeline found at : https://github.com/clairemerot/angsd_pipeline.

## Author Contributions

X Dallaire led the design of the study and the writing of the manuscript, contributed to the production of the data, developed analytical tools, and analyzed the data. R Bouchard, P Hénault, and G Ulmo-Diaz contributed to the production of the data, the design of the study and the writing of the manuscript. E Normandeau and C Mérot contributed some analytical tools, to the design of the study and the writing of the manuscript. JS Moore and L Bernatchez contributed to the design of the study and the writing of the manuscript

## Supplemental Material

**Table S1 :** Proportion masked by RepeatMasker for each repetitive DNA category in the sufficiently part of the genome (>0.75X) and in peaks of coverage (>3X for American Eel, Arctic Char, and Lake Whitefish; >8X for Atlantic Salmon). The χ^2^ statistic for the comparison of proportion is shown.

**Table S2 :** Short Read Archive accession number for Atlantic Salmon sequences

**Table S3 :** Proportion masked by RepeatMasker for each repetitive DNA category in the American Eel, Salvelinus sp., Lake Whitefish, and Altantic Salmon reference genome.

**Table S4 :** Self-syntenic blocks identified by SyMap in the Lake Whitefish, *Salvelinus sp.*, and Atlantic Salmon reference genome.

**Table S5 :** Short Read Archive accession number for the construction of an ancestral genome based of the *Salvelinus sp.* reference genome

**Fig. S1 :** Average depth of coverage by base pair in the Arctic Char, Lake Whitefish (Great Slave Lake then James Bay), Atlantic Salmon, and American Eel datasets.

**Fig. S2 :** Deviant SNPs appear as shared polymorphism between diverged datasets. A) Number of canonical (black) and deviant (red) SNPs in the Lake Whitefish dataset for Great Slave Lake, James Bay, and common SNPs to both datasets. SNPs with different classification depending on the datasets are shown in blue in equal proportion above and below the middle line. B) Relationship between the minor allele frequency of 100,000 SNPs in the two Lake Whitefish datasets. Contours of a kernel density estimation were added for each SNP category to highlight comparison. C) Relationship between F_IS_ of 100,000 SNPs in the two Lake Whitefish datasets.

**Fig. S3 :** Sampling sites for Atlantic Salmon in Norway and the sequencer model used to produce data in Bertolotti et al. 2020

**Fig. S4 :** Rootograms for negative binomial models described in Table 2. The squared root of the frequency for expected (black line with red dots) and observed (hanging histograms) values are compared.

