## Supplemental Figures for "Widespread deviant patterns of heterozygosity in whole-genome sequencing due to autopolyploidy, repeated elements, and duplication"

**Fig. S1** : Average depth of coverage by base pair in the Arctic Char, Lake Whitefish (Great Slave Lake then James Bay), Atlantic Salmon, and American Eel datasets.

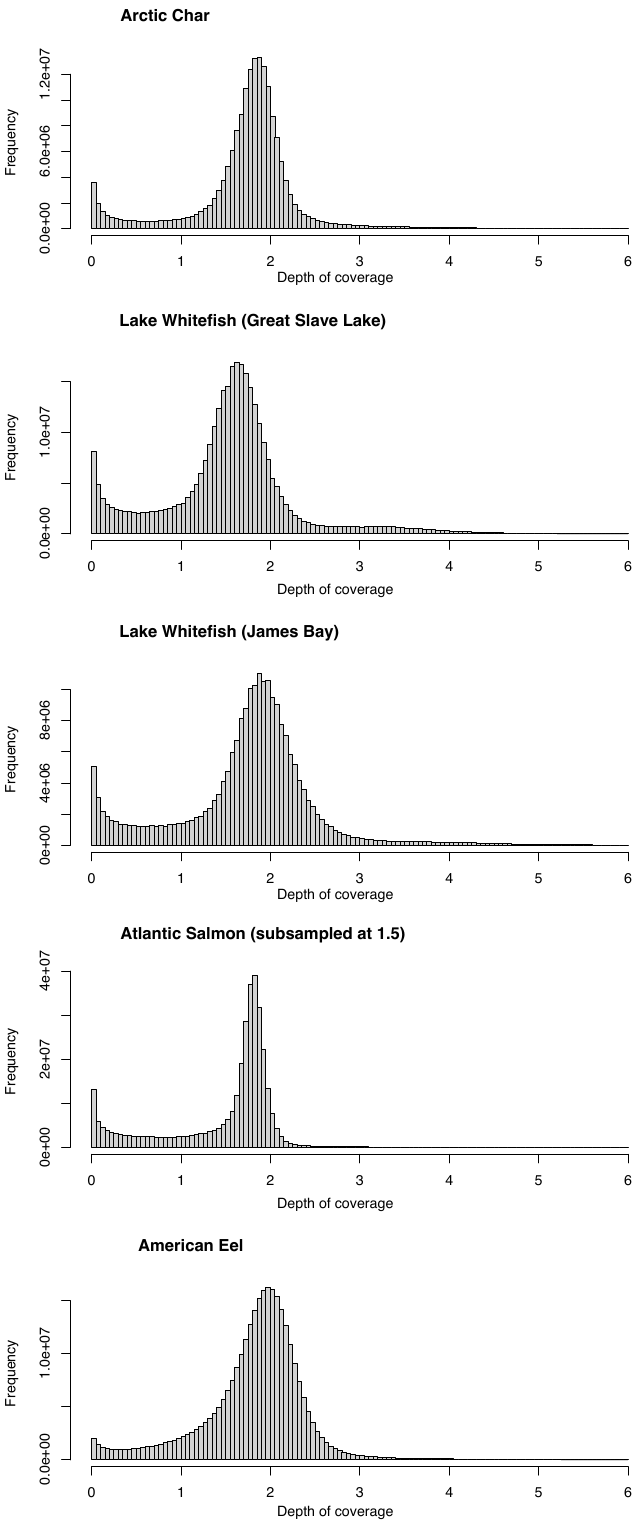

**Fig. S2 :** Deviant SNPs appear as shared polymorphism between diverged datasets. A) Number of canonical (black) and deviant (red) SNPs in the Lake Whitefish dataset for Great Slave Lake, James Bay, and common SNPs to both datasets. SNPs with different classification depending on the datasets are shown in blue in equal proportion above and below the middle line. B) Relationship between the minor allele frequency of 100,000 SNPs in the two Lake Whitefish datasets. Contours of a kernel density estimation were added for each SNP category to highlight comparison. C) Relationship between F_IS_ of 100,000 SNPs in the two Lake Whitefish datasets.

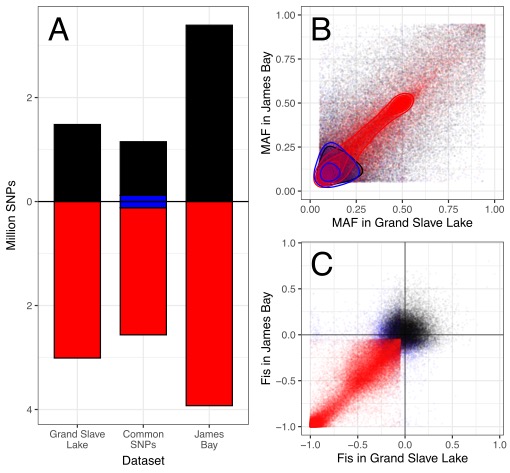

**Fig. S3 :** Sampling sites for Atlantic Salmon in Norway and the sequencer model used to produce data in Bertolotti et al. 2020

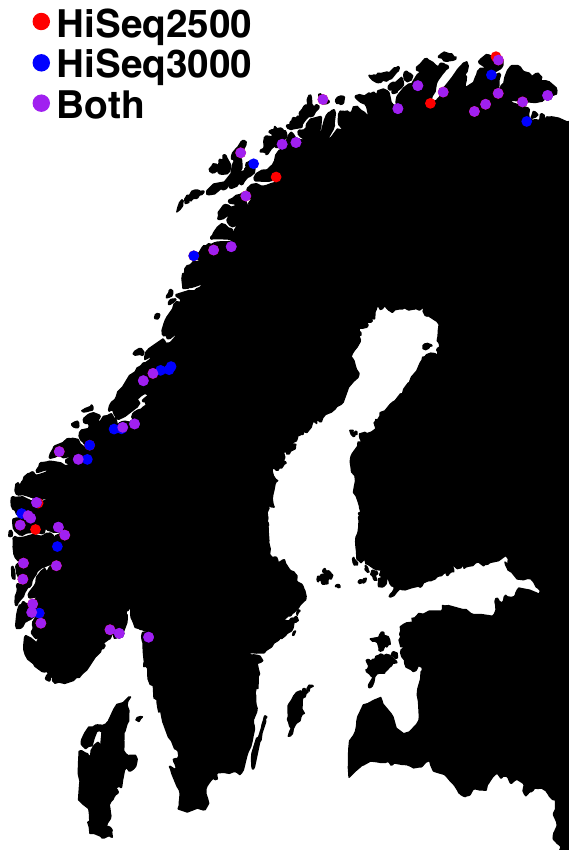

**Fig. S4 :** Rootograms for negative binomial models described in Table 2. The squared root of the frequency for expected (black line with red dots) and observed (hanging histograms) values are compared.

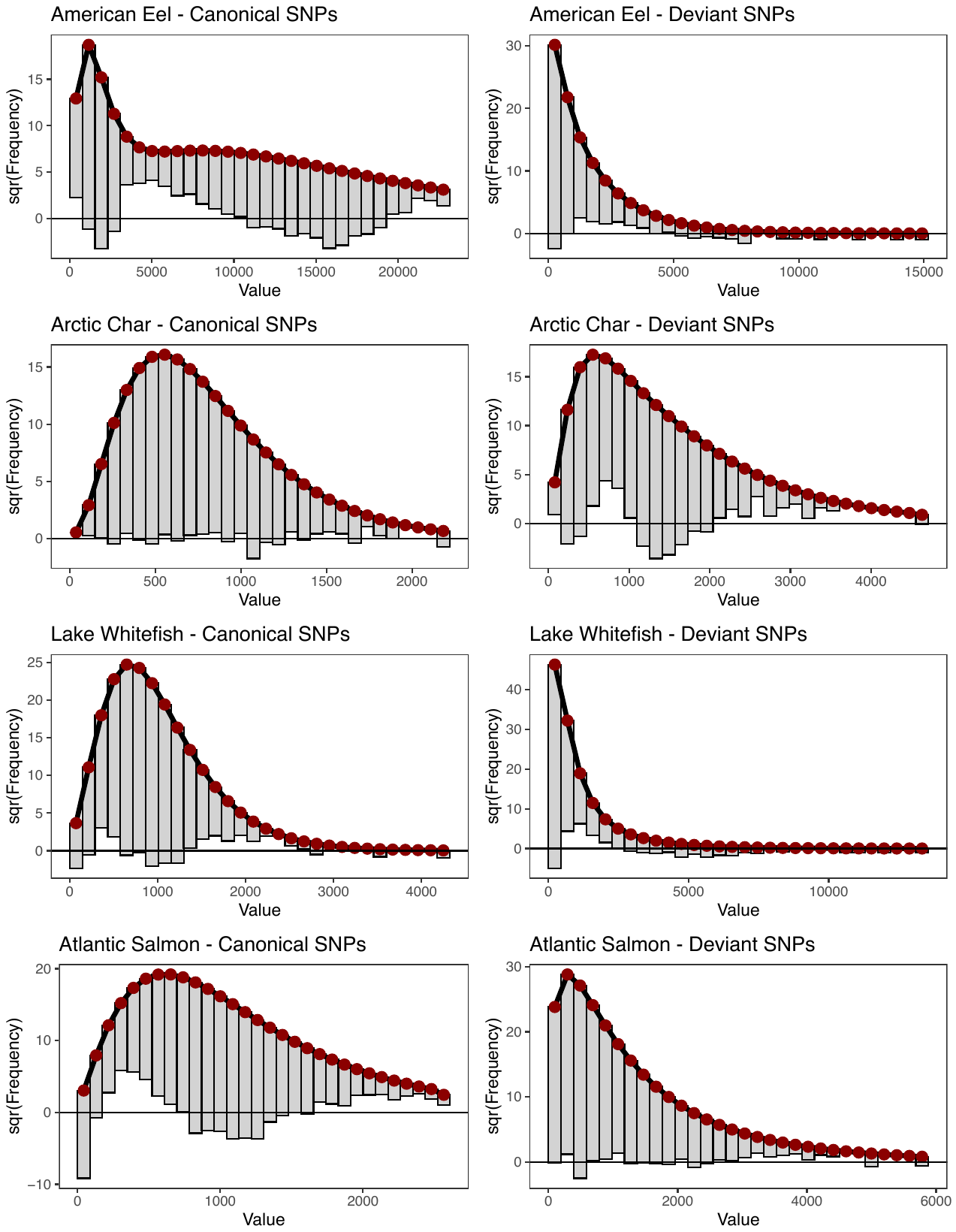
